# Hippocampal collagen as a potential target for post-surgical treatment; effects of whole-body vibration and exercise

**DOI:** 10.1101/2022.11.02.513937

**Authors:** Noa Keijzer, Klaske Oberman, Tamás Oroszi, Csaba Nyakas, Eddy A van der Zee, Regien G Schoemaker

## Abstract

Peripheral surgery may evoke neuroinflammation, associated with neuronal damage and consequently mental health problems. However, anti-inflammatory treatment showed limited therapeutic efficacy. Preservation of neuron integrity during neuroinflammation, by targeting their protective collagen sheet, may provide an alternative strategy. Whole-body vibration (WBV) and exercise combine anti-inflammatory and collagen-increasing effects in the periphery. The present study aimed to explore the therapeutic efficacy of postoperative WBV and exercise on hippocampal neuroinflammation and collagen expression.

Three months old male Wistar rats underwent abdominal surgery. Starting from one day after surgery, rats were submitted to WBV (10 min, once or twice daily, 30 Hz), running exercise (30 min, daily), or pseudo WBV/exercise, for two weeks. Rats were sacrificed and brain tissue was collected and processed for (immuno)histochemistry. Hippocampal microglia activity, total collagen content, and expression of fibrous and non-fibrous collagen subtypes were analysed.

Surgery was associated with increased microglia activity in the CA1 area, which was only partly reversed by the interventions. Surgery specifically reduced total collagen expression in the CA1 area, which was restored by both WBV and exercise. Collagen I was absent in the hippocampal granular layers. The surgery-induced decrease in collagen III expression in the CA1 area was not affected by either WBV or exercise. However, surgery increased collagen III in the CA2 (ns), CA3 and DG. Exercise, and to a lower extent WBV, seemed to (partly) reverse this effect. Collagen IV expression was not altered by surgery, but increased by WBV. No significant effects were observed on collagen VI expression.

WBV as well as exercise restored the surgery-induced declined collagen expression, while partly reversing microglia activation in the CA1 area. Moreover, effects on collagen appeared to be subtype- and region-specific, with overall similar effects of WBV and exercise. Nevertheless, the neuroprotective potential of postoperatively altered brain collagen needs further investigation.

## Introduction

Most people undergo surgery at least once during their life (1). Postoperative complications may develop after surgery, including hippocampal neuroinflammation, which could be associated with cognitive decline and a reduction in patients’ quality of life (2,3). Although aging and previous inflammatory episodes are acknowledged as risk factors, the pathophysiology of this complication is rather unknown (4,5). Most evidence points to a key role for peripheral inflammation and the subsequent induction of neuroinflammation. According to this hypothesis, surgery-induced activation of the native immune system, required for wound healing, can become derailed and is reflected in the brain as neuroinflammation, associated with neuronal dysfunction (6,7). Despite the acknowledged role of neuroinflammation and promising experimental studies (4,6,8,9), the therapeutic potential of anti-inflammatory treatment in the clinic remains poor (10,11).

An alternative strategy may aim at protecting the neurons from the damaging effects of neuroinflammation, rather than, or in combination with, interference of the inflammatory response. A good candidate target could be provided by the net-like structures around the cell body and dendrites of neurons, known as perineuronal nets (PNNs), which protect the neurons against oxidative stress and neurotoxins (12,13). The function of these PNNs in neuronal functions associated with cognitive performance was elegantly reviewed by Wingert and Sorg (14). These specialized substructures of the brain extracellular matrix (ECM) consist of low amounts of fibrous collagens, including type I and III (12). Non-fibrous collagen IV has further been discovered in the vascular basement membrane, another substructure of the brain ECM, that is important for regulating blood-brain barrier (BBB) permeability (15,16). The basal lamina, which comprises a part of the vascular basement membrane, also surrounds PNNs to provide structural support and neuronal protection (17,18). As for collagen VI, a neuroprotective role has previously been described under normal physiological conditions (19), but increased expressions were found upon neuronal injury (20). Since collagen seems essential for preserving brain health, we hypothesized that in addition to the induction of neuroinflammation, certain types of surgery may disrupt the PNNs, potentially leading to a loss of neuronal protection against neuroinflammation-associated neuronal damage. Therapeutic prevention of this surgery-induced loss then may preserve neuronal protection and counteract the potentially detrimental effects of neuroinflammation (21).

Exercise is widely associated with improved physical as well as mental health. It combines anti-inflammatory effects (22) to effects on PNNs (23); increased PNN expression around motor neurons, but declined expression in the hippocampal CA1 area after exercise. Although exercise may provide a beneficial intervention after surgery, not all patients will be capable of performing exercise after their surgical procedure. A less physically straining intervention may be worthwhile. Whole-body vibration (WBV), a specific type of sensory stimulation, in which controlled mechanical vibrations are transmitted to the body and brain via an oscillating platform, may potentially increase brain collagen (24). WBV has been indicated as a passive alternative for conditions in which patients are unable or unmotivated to perform physical exercise, such as during recovery from surgery (25,26). If needed, WBV can even be applied while patients remain in bed and even in an intensive care unit (27). Researchers already demonstrated that WBV enhanced angiogenesis, associated with improved wound healing and lower systemic inflammation, via upregulation of fibrous proteins in diabetic mice (28). Moreover, WBV showed to increase collagen gene expression in the rat patellar tendon (29). Furthermore, WBV promoted an anti-inflammatory response in elderly subjects, similar to that of physical exercise (30), and improved cognitive function in rats (31).

Therefore, since WBV may share the positive effects of exercise but with less physical effort, the aim of the present study was to explore the effects of WBV compared to exercise started shortly after abdominal surgery in rats, regarding neuroinflammation and collagen expression in the hippocampus.

## Materials and methods

### Animals

Three months old male Wistar rats (Janvier, Saint-Isle, France) were group housed (3-4 animals per cage) until the moment of surgery, in climatized rooms (22 ± 2°C; 50 ± 10% humidity, and reversed 12-12 hours (h) light-dark cycle). After the surgical procedure, animals were housed individually. Water and food (standard rodent chow: RMHB/2180, Arie Block BV, Woerden, NL) were available ad libitum. All procedures were approved by the National Competent Authority (CCD) and the local ethical animal welfare committee (IvD) of the University of Groningen, The Netherlands.

### Study design

The present study is a follow-up of the study of Oroszi et al. (32). For details about the animals and the protocol, we kindly refer to this original study. Briefly, three months old male Wistar rats were randomly divided into four groups (n=16 each): 1) non-surgery control, 2) surgery control (+ pseudo WBV/exercise), 3) surgery + WBV and 4) surgery + exercise. Major abdominal surgery consisted of the implantation of a radio telemetry transmitter (33) or standard abdominal surgery; the latter was previously shown to induce neuroinflammation and cognitive decline (34). Non-surgery controls received the same handling, but did not undergo anaesthesia or surgery. Apart from a difference in body weight loss shortly after surgery, no differences in behaviour or neuroinflammation were observed between these two types of surgery, therefore surgery groups were pooled (32). Starting one day after surgery, rats were submitted to either two weeks of WBV, exercise (treadmill running), or pseudo WBV/exercise in the dark (active) phase. On post-operative day 14, animals were deeply anesthetized with pentobarbital (90 mg/kg) and transcardially perfused with cold saline containing Heparin (2 IE/ml). Brains were collected and processed for further histological analyses.

## Interventions

### Whole-body vibration (WBV)

WBV was performed using a low-intensity vibration device with a frequency of 30 Hz and an amplitude of 50-200 microns (MarodyneLiV – Low Intensity Vibration; BTT Health GmbH; Germany). This is in adherence to the newly reported guidelines for WBV studies in animals (35). While on the first post-surgical day, WBV was started with one 10 minutes (min) session, WBV on days 2-6 consisted of two 10 min sessions per day. To avoid interference with behavioral testing during the second post-surgical week, sessions were reduced to once daily and performed after the behavioral testing procedures. After the behavioral test week, animals returned to WBV sessions twice daily until sacrifice.

### Running exercise

The exercise was performed by running on a treadmill (Home-made, University of Groningen, The Netherlands) for 30 min daily, starting one day after the surgery. Training started at a low speed (5-10 m/min) and was gradually increased to the aimed speed of 18 m/min, which would reflect approximately 65% of the maximum oxygen uptake (36).

### Pseudo WBV/Running exercise

Pseudo-treated rats were subjected to a combined pseudo WBV and pseudo running exercise treatment in order to serve as a control for both interventions, based on our previous experiments. This consisted of daily alternating twice 10 min (once 10 min during the testing phase) on the vibration plate without vibration or 30 min on the turned-off treadmill.

### Histology

#### Tissue preparation

Brains were collected and post-fixated in 4% paraformaldehyde solution for two days, and washed with 0.01M phosphate-buffered saline (PBS) for three consecutive days. For cryopreservation, brains were dehydrated for one day with a 30% sucrose solution before freezing with liquid nitrogen, and stored at -80 °C. Coronal sections (25 µm) of the dorsal hippocampus were freshly cut and collected directly on Superfrost™ Plus Microscope Slides, or stored free-floating in 0.01M PBS + 0.1% Natrium Azide at 4°C for later use.

#### Ionized binding adaptor protein-1 (IBA-1) immunostaining

Microglia activity was used as a measure of hippocampal neuroinflammation, as previously described elsewhere (37). Microglia were immunohistochemically visualized by incubating the free-floating dorsal hippocampal sections with a rabbit-anti-ionized binding adaptor protein-1 primary antibody (IBA-1; Wako, Neuss, Germany, 1:2500) in 2% bovine serum albumin and 0.1% Triton X-100 for three days at 4°C, followed by incubation with a goat anti-rabbit secondary antibody (Jackson, Wet Grove, USA, 1:500) for 1 h at room temperature. Individual images of the cornu ammonis (CA)1, CA2, CA3, CA4, dentate gyrus (DG) inner blade (DGib), DG outer blade (DGob), and Hilus areas were taken at 200x magnification. Image-Pro Plus 6.0 (Media Cybernetics, Rockville, USA) was used to obtain microglia morphology parameters. Microglia were analysed regarding the density (number per high power field), coverage (%), cell size (pixels), cell body size (pixels), and dendritic processes size (pixels). Microglia activity was calculated as cell body size per total cell size (%) (37). Values of the Dgib and DGob regions were averaged resulting in the DG area value.

#### Sirius Red/Fast Green histochemical staining

Freshly cut dorsal hippocampal sections were collected on Superfrost™ Plus Microscope Slides and allowed to dry before staining of total collagen as described in detail elsewhere (34). Briefly, dried sections were incubated for 30 min with 0.1% Sirius red (Sigma-Aldrich), followed by 30 min incubation with 0.1% Fast green (Sigma-Aldrich), both dissolved in saturated picric acid. Staining steps were followed by rinsing in 0.01M hydrochloric acid for 1 min (2x), tap water for 1 min (2x), and in demi water for 1 min (2x) and cover-slipped. Stained sections were automatically scanned using a Nanozoomer 2.0-HT digital slide scanner (Hamamatsu, Japan). Individual images were taken of the CA1a, CA1b, CA2, CA3, CA4, DGib and DGob at 40x magnification. The area of total collagen in a predetermined area of interest in the granular layer was quantified and expressed as a percentage coverage (Image-Pro Plus 6.0, Media Cybernetics, Rockville, USA). Values of the CA1a/CA1b and DGib/DGob regions were averaged resulting in the CA1 and DG area, respectively.

#### Collagen I, III, IV, and VI immunostaining

Free-floating dorsal hippocampal sections were incubated for 5 min in demi water at 37 °C, followed by treatment with 0.5 mg pepsin/ml (Roche) for 18 min at 37 °C. Sections were then treated for 30 min with 0.3% H_2_O_2_ and incubated with rabbit anti-collagen I (Abcam, ab270993, 1:2000), mouse anti-collagen III (Abcam, ab6310, 1:2000), rabbit anti-collagen IV (Abcam, ab6586, 1:2500) or rabbit anti-collagen VI (Novus Biologicals, NB120-6588, 1:100) in 1% bovine serum albumin and 0.1% Triton X-100 for three days at 4°C. Next, sections were incubated for 2 h with a goat-anti-rabbit (Jackson ImmunoResearch, 1:500) or goat-anti-mouse (Jackson ImmunoResearch, 1:500) secondary antibody, followed by a 1 h incubation with avidin-biotin-peroxidase complex (Vectastain® Elite ABC-HRP Kit, 1:500). Labelling was visualized using a 3,3’-Diaminobenzidine solution (Sigma-Aldrich) activated by 0.1% H_2_O_2._ Sections were mounted onto microscope slides using a 1% gelatin/0.05% aluin solution and submitted to a dehydration process (70% ethanol for 5 min, 100% ethanol for 5 min (2x), 70% ethanol/30% xylene for 5 min, 30% ethanol/70% xylene for 5 min and 100% xylene for 5 min (3x) and covered. All dilutions were made in 0.01M PBS and all sections were washed 3-6 times with 0.01M PBS between staining steps. Negative control staining was performed by replacing the primary antibodies with 0.01M PBS. For positive control, positive staining of blood vessels and meninges was used. Collagen III expression was scored manually as presence/absence due to the weak staining signal in the CA1a, CA1b, CA2, CA3, CA4, DGib and DGob regions and converted in percentages (%). For collagen IV and VI, stained sections were scanned using the Nanozoomer 2.0-HT digital slide scanner and individual images of the CA1a, CA1b, CA2, CA3, CA4, DGib, and DGob were taken at 40x magnification. The optical density (OD) was measured (ImageJ 2.1.0, USA) and corrected for the background staining. The average OD in each hippocampal area was taken as a measure for collagen IV and VI expressions. For all stainings, values of the CA1a/CA1b and DGib/DGob regions were averaged resulting in the CA1 and DG area, respectively.

### Statistical analysis

All statistical analyses were performed using IBM SPSS Statistical Software version 27.0.1 (IBM SPSS Statistics, Armonk, NY). Data outside twice the standard deviation of its group were regarded as outliers and omitted from further statistical analyses (max 2 per group). A Shapiro-Wilk test was performed to test for normality and a Levene’s test to verify the homogeneity of variances. A one-way ANOVA analysis followed by a Least Significant Difference (LSD) post hoc test was done for normally distributed data with an equal variance, to discover surgery and intervention-related differences. Otherwise, a Kruskal Wallis test and Mann-Whitney test were performed to reveal group differences. Differences among groups were considered statistically significant at p < 0.05 (*). Relevant tendencies (one-way ANOVA: p < 0.15, LSD post hoc: p < 0.05 or p < 0.01) were also mentioned (# or ##). Figures were prepared using GraphPad Prism (version 5.00 for Windows, GraphPad Software, San Diego, California, USA) and data in the figures and tables are expressed as mean ± SEM per group.

## Results

### General

Before surgery, rats weighed on average 369 ± 4 grams without differences between the experimental groups. From the initial 64 rats (16 per group), 3 rats died around the surgical procedure, resulting in the following experimental groups: 1) non-surgery control (n=14), 2) surgery control (+ pseudo WBV/exercise) (n=16), 3) surgery + WBV (n=16) and 4) surgery + exercise (n=15). All surgery rats lost significant weight, but without the effects of interventions (data not shown).

### Effects on hippocampal microglia activity and morphology

To assess the effects on hippocampal neuroinflammation, an IBA-1 staining was performed as shown in Fig 1. Fig 1A displays the photomicrographs of the CA1 region of the four experimental groups. Results of microglia activity are shown in Fig 1B for the CA1, and Figs 1C-G for the other hippocampal areas. Analyses of differences between groups showed a tendency for microglia activation in the CA1 area after surgery (p = 0.131; post hoc analyses p < 0.020) (Fig 1B), but not in the other hippocampal areas. WBV and exercise interventions only partly affected hippocampal microglia activity in this area, as results did not appear significantly different from non-surgery. No significant differences were observed in other hippocampal areas (Figs 1C-G). Underlying measurements of microglial morphology are presented in Table 1. These data support the absence of effects on microglia morphology by surgery or interventions in hippocampal areas, other than CA1.

**Table 1.**
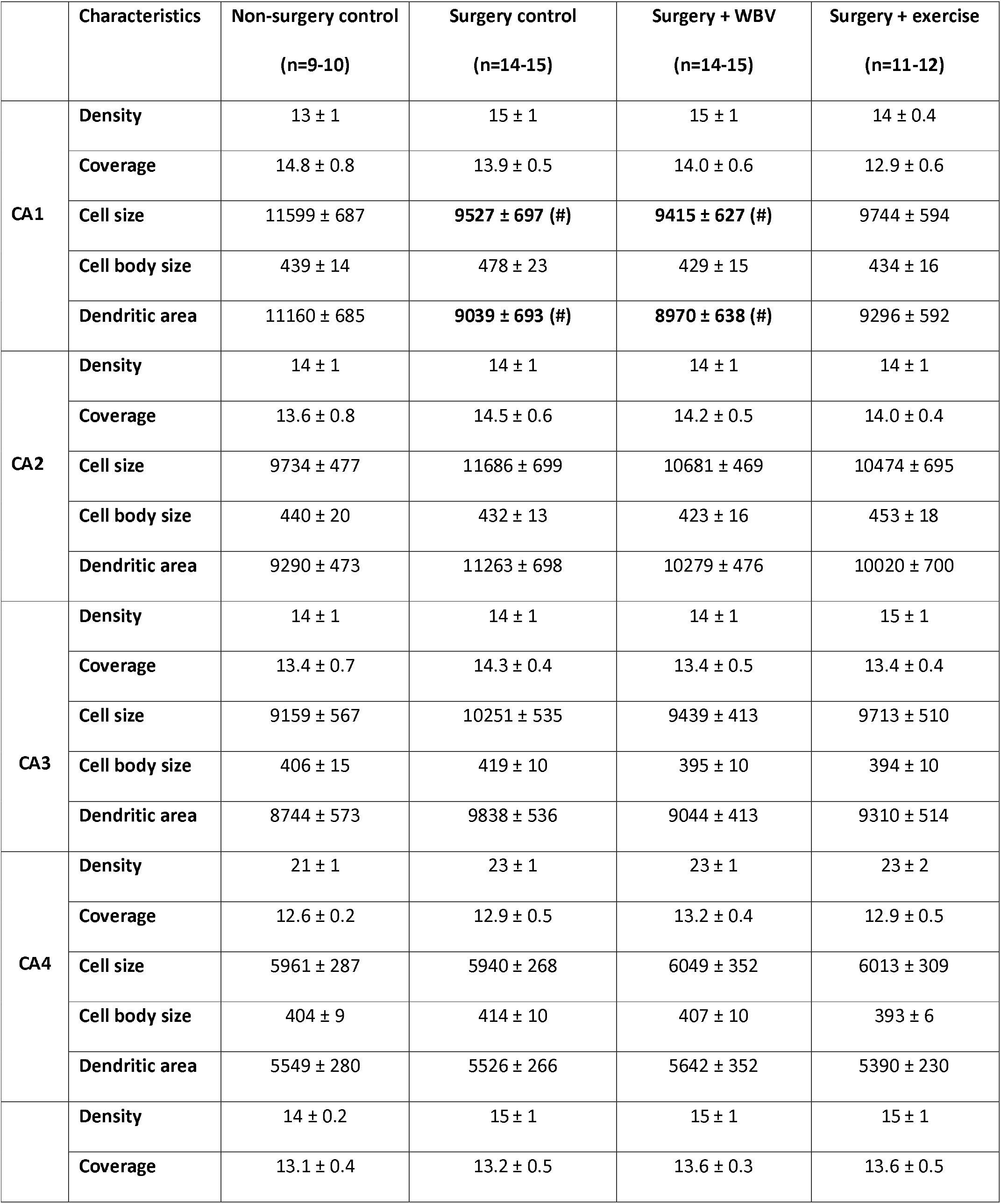
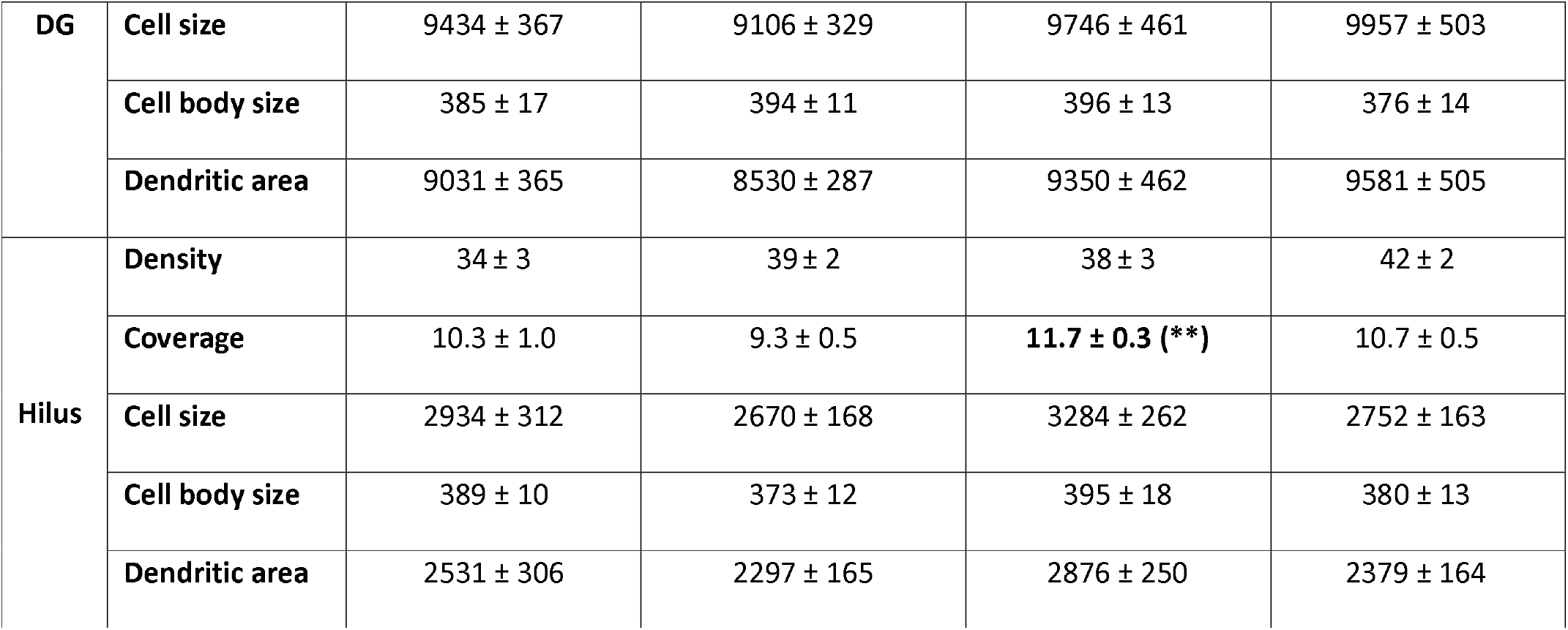
Microglia morphology characteristics in the different rat hippocampal areas; CA1, CA2, CA3, CA4 and DG (mean ± SEM). The density (number per high power field), coverage (%), cell size (pixels), cell body size (pixels) and dendritic processes size (pixels) of the microglia images are shown. **: significant (p < 0.01) difference compared to surgery control. #: relevant tendency (one-way ANOVA: p < 0.150, LSD post hoc: p < 0.05). Differences and relevant tendencies are presented bold. CA: Cornu Ammonis, DG: Dentate Gyrus.

**Fig 1.**
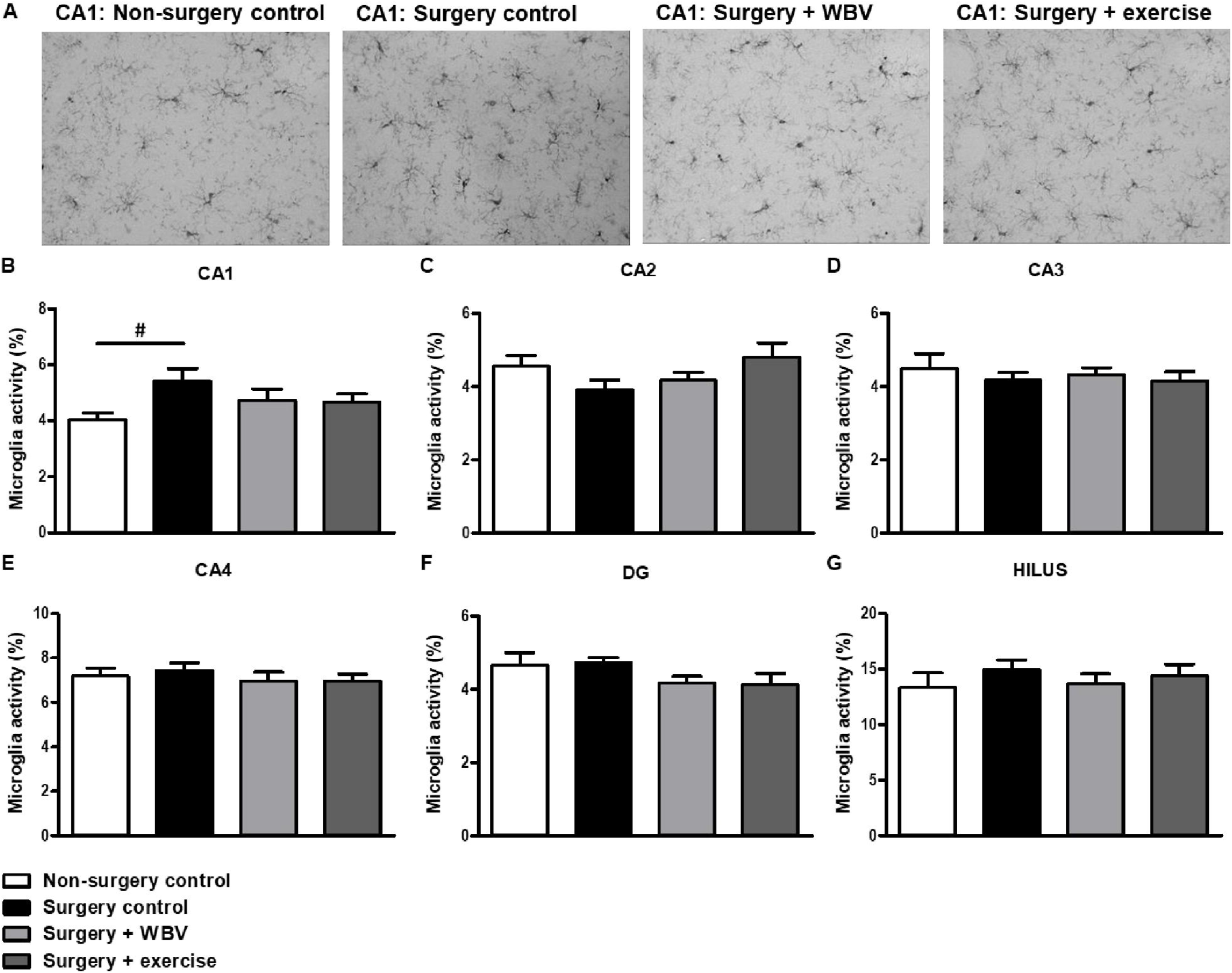
Microglia activity in rat hippocampal areas (mean ± SEM). A: Representative images of microglia staining in the hippocampal CA1 area in the different experimental groups. B-G: Microglia activity, measured as the ratio between cell body size and total cell size in the hippocampal CA1, CA2, CA3, CA4, DG and Hilus region. #: relevant tendency (one-way ANOVA: p < 0.150, LSD post hoc: p < 0.05) between indicated groups. Non-surgery control (n = 9-10), surgery control (+ pseudo WBV/exercise) (n = 14-15), surgery + WBV (n = 14-15) and surgery + exercise (n = 11-12). CA: Cornu Ammonis, DG: Dentate Gyrus, WBV: Whole-body vibration.

### Effects on hippocampal total collagen

The effect of abdominal surgery followed by two weeks of pseudo WBV/exercise, WBV or exercise intervention on total hippocampal collagen was determined with a Sirius red/Fast green histochemical staining, as shown in Fig 2. The upper left panel shows a representative photomicrograph of the hippocampus, indicating positive Sirius red staining of blood vessels (arrows) and meninges, and a Sirius red positive signal in the granular layers of the hippocampus (Fig 2A).

**Fig 2.**
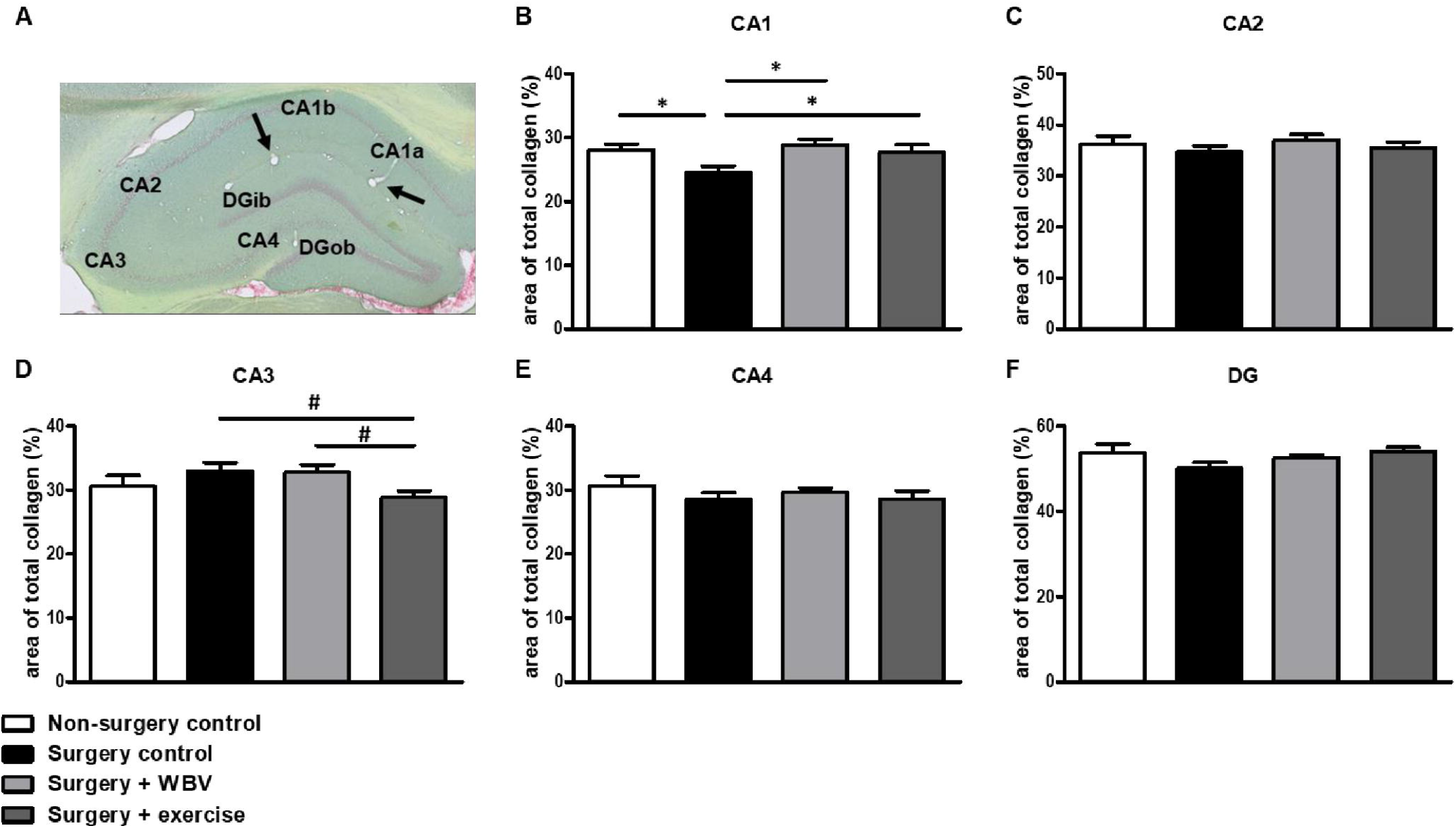
Representative photomicrograph of Sirius red/Fast green stained hippocampus and identified areas, as well as quantification of hippocampal total collagen (mean ± SEM). A: Sirius Red/Fast Green staining was performed to visualize hippocampal total collagen in the CA1, CA2, CA3, CA4 and DG areas. Positive-stained blood vessels are indicated with arrows. The area of total collagen (%) was quantified in predetermined areas of interest (B-F). *: significant (p < 0.05) difference between indicated groups. #: relevant tendency (one-way ANOVA: p < 0.150, LSD post hoc: p < 0.05) between indicated groups. Non-surgery control (n = 12-13), surgery control (+ pseudo WBV/exercise) (n = 16), surgery + WBV (n = 13-15) and surgery + exercise (n = 12-13). CA: Cornu Ammonis, DG: Dentate Gyrus, WBV: Whole-body vibration.

Collagen was mainly observed around the neuronal cell bodies in the hippocampal granular layer. Measurements of the percentage of total collagen expression per hippocampal area are presented in the other panels (Figs 2B-F). Surgery significantly reduced collagen expression in the CA1 area (Fig 2B), but not in other areas (Figs 2C-F). Postoperative WBV and exercise both significantly recovered this declined collagen expression in the CA1. In the CA3 area, a tendency towards reduced total collagen expression was observed after exercise intervention, but not after WBV, when compared to surgery controls (Fig 2D).

### Effects on hippocampal fibrillar collagen I and III

Immunohistochemical staining of hippocampal collagen I demonstrated a clear presence of fibrillar collagen I in the meninges and blood vessels. However, no staining was observed in or around the neurons in the hippocampal granular layers (data not shown). Fibrillar collagen III, although clearly present in blood vessels (arrows) and meninges, showed low expression around the neurons in the different granular layer regions, hampering accurate quantification (Fig 3A). Therefore, the presence/absence of collagen III expression was scored per hippocampal area (Figs 3B-F). In the CA1 area, collagen III was significantly less present after surgery compared to non-surgery controls, but this was not affected by either intervention. In contrast, surgery induced a significant rise in collagen III presence in the CA3 and DG areas. In the CA2 area, a similar, but not significant increase, was observed, which was completely normalized by WBV. Moreover, two weeks of postoperative running exercise significantly decreased collagen III presence in the CA2 and CA3 areas.

**Fig 3.**
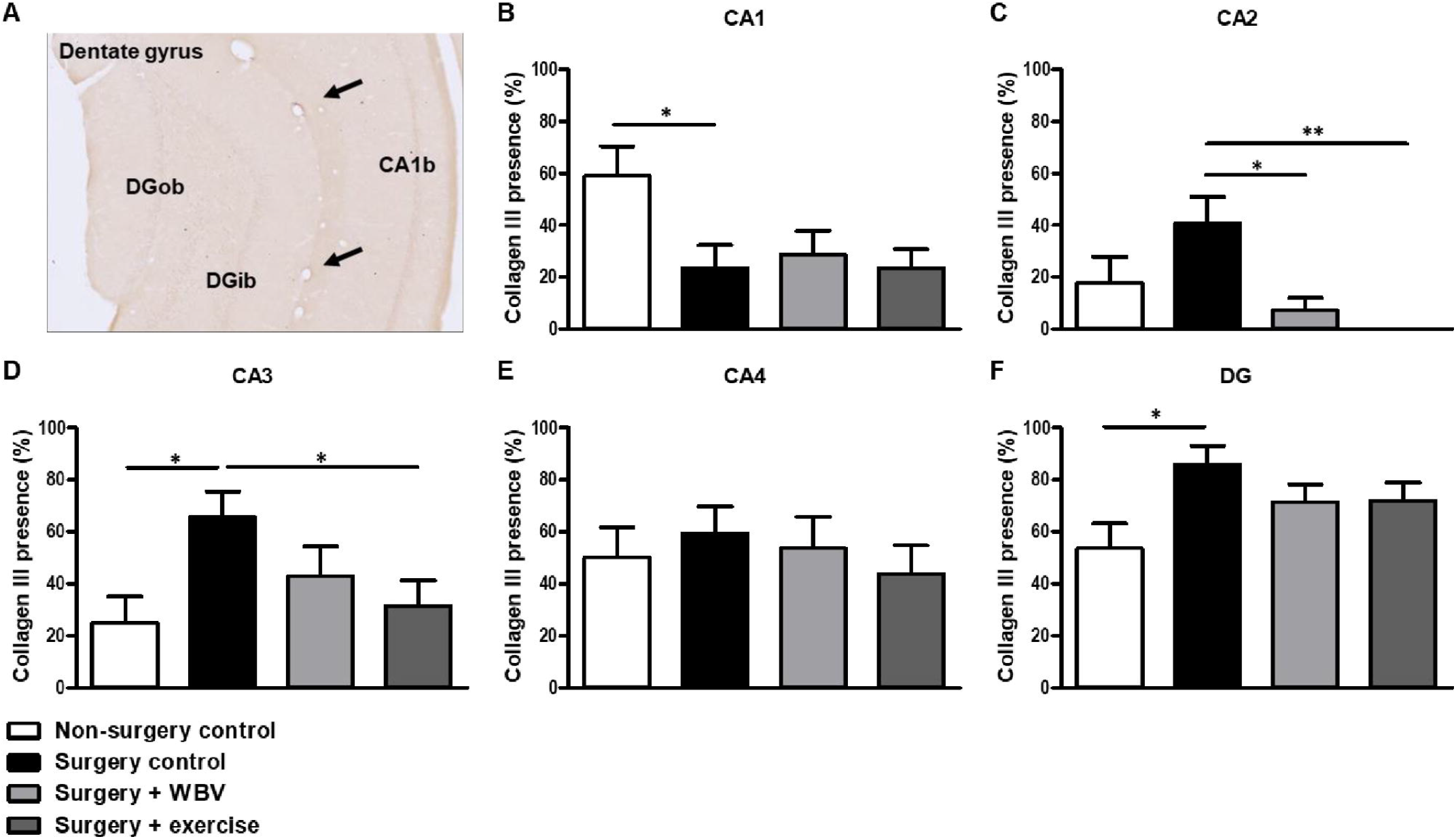
Representative photomicrograph of a hippocampus stained for collagen III and quantification of hippocampal collagen III presence (mean ± SEM). A: Representative image of immunohistochemical collagen III staining in the hippocampus. Positive-stained blood vessels are indicated with arrows. B-F: The presence of collagen III was manually scored in the CA1, CA2, CA3, CA4 and DG granular layer. *: significant (p < 0.05) or **: significant (p < 0.01) difference between indicated groups. Non-surgery control (n = 14), surgery control (+ pseudo WBV/exercise) (n = 16), surgery + WBV (n = 16) and surgery + exercise (n = 15). CA: Cornu Ammonis, DG: Dentate Gyrus, WBV: Whole-body vibration.

### Effects on hippocampal non-fibrillar collagen IV and VI

Effects of the abdominal surgery or WBV and exercise interventions on non-fibrillar collagen subtypes were obtained from measurements of collagen IV and VI expressions. Hippocampal collagen IV expression is illustrated in Fig 4A and quantified in Figs 4B-F. Responses seemed uniform in the different hippocampal areas; defined as no significant effects of abdominal surgery on hippocampal collagen IV expression while postoperative WBV, but not running exercise, seemed to increase collagen IV expression in all regions compared to non-surgery controls and surgery controls. However, these findings only reached statistical significance in the CA2 and CA3 areas.

**Fig 4.**
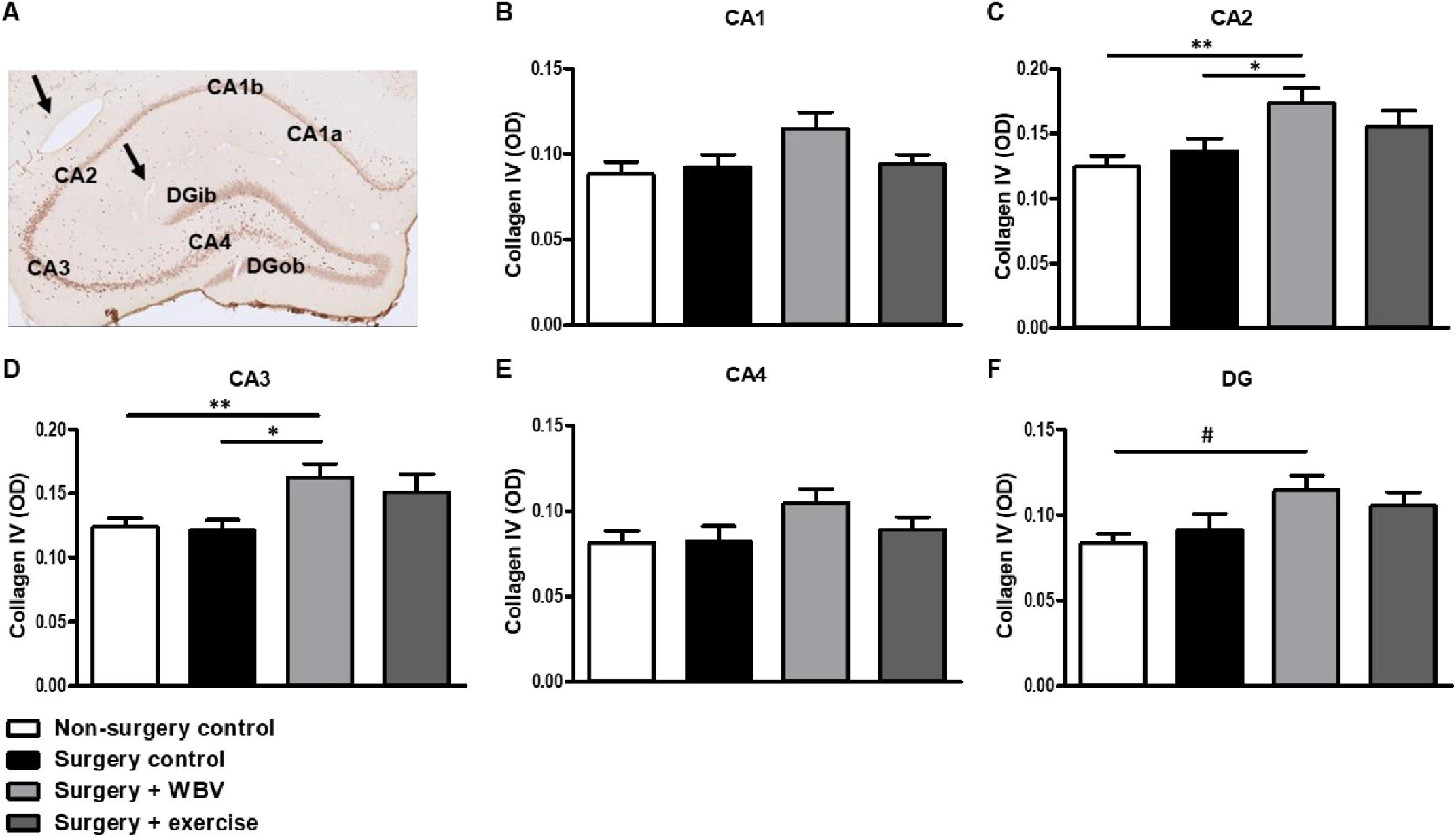
Representative photomicrograph of a hippocampus stained for collagen IV and quantification of hippocampal collagen IV expression (mean ± SEM). A: An immunohistochemical collagen IV staining showed clear presence of collagen IV in the CA1, CA2, CA3, CA4 and DG granular layer. Positive-stained blood vessels are indicated with arrows. B-F: Collagen IV expressions were quantified by measuring the OD in the individual hippocampal areas. *: significant (p < 0.05) or **: significant (p < 0.01) between indicated groups. #: relevant tendency (one-way ANOVA: p < 0.150, LSD post hoc: p < 0.05). Non-surgery control (n = 13-14), surgery control (+ pseudo WBV/exercise) (n = 15-16), surgery + WBV (n = 15-16) and surgery + exercise (n = 15-16). CA: Cornu Ammonis, DG: Dentate Gyrus, OD: Optical Density, WBV: Whole-body vibration.

Similar to collagen IV, no effect of surgery was observed on collagen VI expression in the hippocampal areas (Fig 5). Moreover, neither intervention significantly affected collagen VI expression, although a tendency towards an increase was observed after exercise in the CA1 area (Fig 5B).

**Fig 5.**
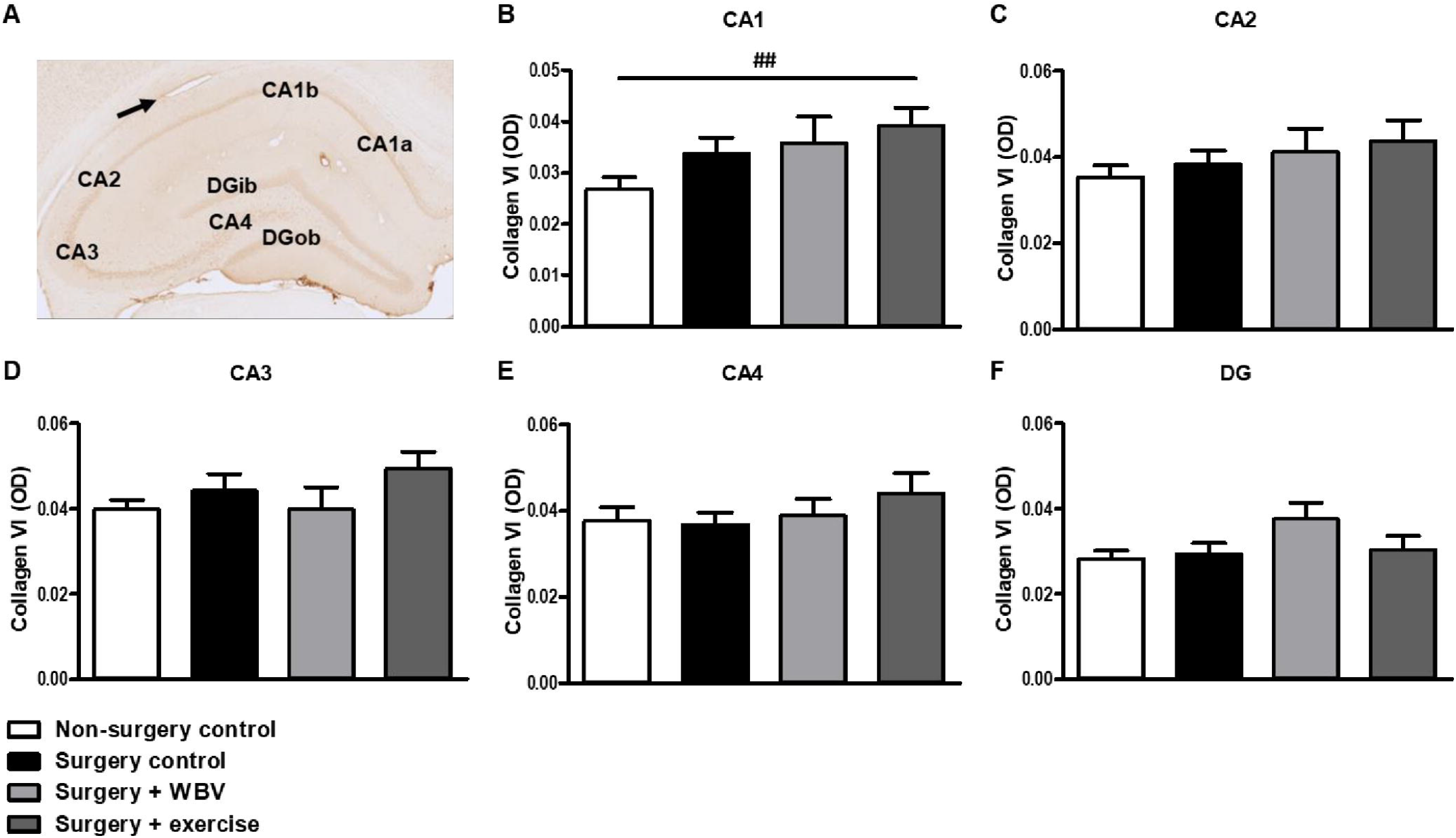
Representative photomicrograph of a hippocampus stained for collagen VI and quantification of hippocampal collagen VI expression (mean ± SEM). A: An immunohistochemical collagen VI staining showed clear presence of collagen VI in the CA1, CA2, CA3, CA4 and DG areas. A positive-stained blood vessel is indicated with an arrow. B-F: Quantification of collagen VI expression by measuring the OD in the individual hippocampal areas. ##: relevant tendency (one-way ANOVA: p < 0.150, LSD post hoc: p < 0.01). Non-surgery control (n = 8-10), surgery control (+ pseudo WBV/exercise) (9-11), surgery + WBV (n = 5-8), surgery + exercise (n = 9-10). CA: Cornu Ammonis, DG: Dentate Gyrus, OD: Optical Density, WBV: Whole-body vibration.

## Discussion

### General

Post-surgical mental complications may be attributed to neuronal damage, associated with neuroinflammation. In addition to inhibiting neuroinflammation, neuroprotection after surgery may be obtained by preserving the protective collagen sheets around neurons; the PNNs. Since WBV and exercise share anti-inflammatory and collagen-increasing effects in the peripheral part of the body, the aim of the present study was to investigate the effects of WBV and exercise on the hippocampus after surgery. Indeed, in addition to induction of neuroinflammation, surgery decreased total collagen expression in the CA1 area of the hippocampus. Whereas two weeks of postoperative WBV or exercise only partly reversed hippocampal neuroinflammation, both interventions significantly prevented the surgery-induced decrease in total hippocampal collagen. Other hippocampal areas were not significantly affected. A more detailed investigation of the expression of collagen subtypes revealed a region-specific and mixed response of fibrillar and nonfibrillar collagen subtypes. Hence, our data indicate that hippocampal collagen can be quantified and seemed sensitive to the effects of surgery as well as to the effects of WBV and exercise. The contribution of the different subtypes of collagen to neuroprotection as well as more general hippocampal functions are subjects for follow-up studies.

### WBV and exercise

The aim of the present study was to compare WBV to exercise early after surgery. WBV and exercise share systemic effects, including increased muscle strength (38–40), and improved wound healing, as well as effects on the brain, such as increased neurotrophic factors (41,42), and cognitive improvement (31,43,44). As reviewed by Alam and coworkers (45), WBV is regarded as a neuromuscular training method to be used as an alternative to conventional training and therapy. WBV was reported to reduce brain damage and brain inflammatory markers, with increased brain-derived neurotrophic factor and improved functional activity after transient brain ischemia in middle-aged female rats (41). These studies support the potential of WBV as an alternative for exercise in the early phases after surgery.

### Effects on neuroinflammation

Effects on neuroinflammation were determined based on changes in microglia morphology; microglia activity was calculated as cell body to cell size ratio (37). Microglia activation is associated with increased cell body size at the expense of processes (46,47). Our previous studies indicated that abdominal surgery in young healthy male rats would increase microglia activity, predominantly in the CA1 area of the hippocampus (4,34,48). In the present study, CA1 microglia activation after surgery did not reach statistical significance, however, an increase of over 30% is comparable to our previous findings. Although previous research indicated that WBV and exercise share anti-inflammatory effects (30), in the present study, both interventions only partly restored CA1 microglia activity. This may relate to the duration and/or intensity of the interventions.

### Effects on hippocampal collagen

#### Total collagen

In the brain, collagen is essential for the stabilization of brain ECM structures, such as PNNs and the vascular basement membrane together with the basal lamina, to provide protection of neurons. These PNNs mainly cover the cell body and dendrites and play an important role in learning, memory and information processing in health as well as disease (12). More specifically, it may affect synaptic morphology and function. Loss of PNNs is often seen in neurodegenerative diseases, as reviewed by Bonneh-Barkay and Wiley (49). Moreover, PNNs around hippocampal interneurons can resist destruction by activated microglia (21). In the hippocampus, particularly the CA1, CA3, and DG regions are essential for memory encoding, retrieval of complete memories from partial information and spatial pattern separation (50,51). Overall, the CA1 area comprises the primary output of the hippocampus to other brain regions (52). In our previous studies (34,48), surgery-induced hippocampal neuroinflammation and decreased neurogenesis were associated with a hippocampus-associated decline in short-term and long-term spatial memory. As anticipated, in our present study, total collagen expression was found to be decreased after surgery in the CA1 area, as one of the early hippocampal areas affected by surgery and inflammation (48), but not in the other hippocampal areas. This decreased collagen in the CA1 may then be regarded as a surgery-associated loss of neuroprotection (12). WBV and exercise specifically restored total collagen in the hippocampal CA1 region, which could be regarded as preservation of neuroprotection. On the other hand, in the hippocampal CA2 area, PNNs have been described to restrict synaptic plasticity (53). In the present study, exercise seemed to decrease collagen in the CA3 area. If collagen levels may reflect synaptic plasticity, as described for the CA2 (53), the reduction of collagen by exercise may then be speculated as an improvement of synaptic plasticity. In our previous study, the effects of surgery and reversal by WBV and exercise were observed on cognitive flexibility, rather than on short-term and long-term memory (32). If indeed total hippocampal collagen may reflect aspects of the PNNs function, such as restriction of plasticity during adulthood, this may have played a role in the effects on cognitive flexibility (14). Additional behavioral tests, for instance on pattern completion as a test for CA3 function (54), could shed more light on potential associations between brain collagen, neuronal function and consequently behavior (55).

#### Fibrillar collagen I and III

In mammals, the fibrous collagens I, II, III, V, and XI are mainly present, with collagen I and III making up 95% of total body collagen (56). Whereas both types are important for structural support of the ECM, collagen III is mainly associated with injury and wound healing (57). Under healthy physiological conditions, collagen I and III can be found in a 4:1 ratio, which shifts towards an increase in collagen III during pathological states characterized by tissue damage in the body (58). In the brain, still little is known about collagen presence and function due to the long-standing belief that collagen is absent in the mammalian central nervous system (CNS) (59). However, Hubert et al. have provided a review on the function of several collagen types during the development of the CNS, but also during pathological states in both animals and mammalians (60). Although collagen I is the most abundant protein in the mammalian body (56) and turned out to be present in the brain blood vessels and meninges, collagen I appeared virtually absent in the hippocampus in the present study. Similar to total collagen, a declined collagen III presence was observed in the CA1 area after surgery. Based on the responses to injury in the body (58), an increased rather than decreased collagen III expression after surgery may be anticipated. However, an elevated presence was observed in the hippocampal CA2, CA3 and DG regions after surgery. If indeed collagen III expression reflects tissue damage (58), this may suggest that these latter areas were more affected by neuronal injury. The hippocampal CA2 region is essential for social memory (61) and Zhang et al. (62) recently demonstrated that elderly patients who underwent anaesthesia and cardiac surgery developed social cognitive dysfunction, indicating that social memory may be affected. For the hippocampal CA3 region, the most important functions are the rapid encoding of memory (63) and retrieval of complete memories (64). Postoperative WBV and physical exercise both reversed the surgery-induced increased presence of collagen III in the CA2 and CA3 region, potentially suggesting repair and a neuroprotective effect. With WBV being less physically demanding than exercise, WBV may provide a relevant alternative for patients who are not capable of performing exercise shortly after surgery.

#### Non-fibrillar collagen IV and VI

In contrast to fibrous collagens, non-fibrous collagens are expressed at much lower levels in the body. Their main function is to adjust structural characteristics, such as the shape and fibre thickness of collagen I, or to connect fibre groups to each other or surrounding tissue (65). Given that collagen IV is essential for BBB integrity (15,16), may indicate that a decrease in collagen IV permits BBB leakage. In contrast to a study from Cao et al. in which four hours isoflurane anaesthesia decreased collagen IV in brain blood vessels leading to BBB disruptions (66), isoflurane anaesthesia and abdominal surgery did not decrease hippocampal collagen IV expressions in the present study. However, Cao et al. measured collagen IV expression immediately after anaesthesia, while in our study, measurements were performed two weeks after anaesthesia and surgery. Therefore, collagen IV levels might already be recovered and not contribute to any further long-term BBB leakage after surgery as seen previously (67). Nevertheless, our finding that postoperative WBV increased hippocampal CA2 and CA3 collagen IV levels and a relevant tendency was observed in the DG, could be regarded as a potential neuroprotective effect of WBV. In addition to its essential role in maintaining BBB integrity, the vascular basement membrane is an important contributor to the development of brain blood vessels (68). Cavaglia et al. previously demonstrated that the rat microvessel density was higher in the hippocampal CA3 region compared to the CA1 region (69), indicating hippocampal area-specific differences in capillary density, which may be reflected in area-specific differences in collagen IV expression responses to WBV. As for exercise, Davis et al. demonstrated that running exercise (30 min daily for a total of 3 weeks) increased collagen IV expression and reduced its loss after stroke in rat cortex and striatum basal lamina, hence, indicating improved BBB integrity and basal lamina function (70). Accordingly, in the present study, exercise may have induced a slight, but not significant, increase in CA3 collagen IV expression. Longer exercise intervention may be needed to induce a significant effect. Collagen VI, which was previously demonstrated to have a neuroprotective role under physiological conditions (19), and expression was shown to increase upon neuronal injury (20), showed no significant differences in expression in the different hippocampal areas after abdominal surgery or after either intervention. These findings may implicate that the rats did not display severe hippocampal neuronal injury two weeks after surgery, as was hypothesized. Accordingly, microglia activation was only observed in the CA1 area, and in general agreement with our previous findings (4,34,48).

## Conclusions

Results indicate that surgery, in addition to the induction of microglia activation, may decline collagen expression in the hippocampus (CA1). This neuroinflammation combined with a loss of neuroprotection may facilitate neuronal damage. Although only partly restoring neuroinflammation, both exercise as well as WBV normalized collagen expression after surgery. The functional consequences of these area-specific and collagen subtype-specific effects for neuroprotection and subsequent behaviour responses need further investigation. Nevertheless, since WBV shared effects seen with exercise, WBV may provide a promising intervention for patients not capable or motivated to perform exercise, either as replacement or step-up to exercise.

## Supporting information

cover letter

## Acknowledgments

We thank Kunja Slopsema for her technical support and master student Eleni Gkotski for her help in the microglia analysis. Erjan Hilbrands and Jacko Duker from the Department of Pathology at the University Medical Center Groningen are also greatly appreciated for digital scanning of the stained sections.

