## Supplementary material for "Hippocampal collagen as a potential target for post-surgical treatment; effects of whole-body vibration and exercise": cover letter

| Faculty of Science and Engineering |  | behavioral and cognitive neurosciences |
| --- | --- | --- |

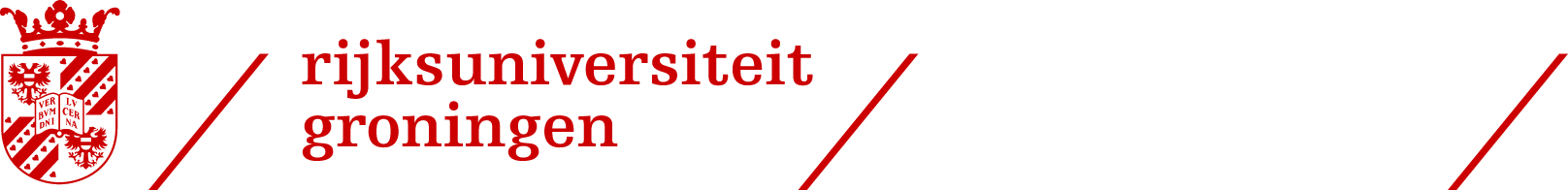

| Editor in Chief,  Frontiers in Aging Neuroscience | |  | Nijenborgh 7 | |
| --- | --- | --- | --- | --- |
|  |  |  | Department of Molecular Neurobiology | |
|  |  |  | Groningen, 9747 AG  The Netherlands | |
|  |  |  | Phone: | +31 (050) 363 2299 |
|  |  |  | e-mail: | |
| Date November 1, 2022 |  |  |  |  |
| Subject: Cover letter Title: **Hippocampal collagen as a potential target for post-surgical treatment; effects of whole-body vibration and exercise** | | | | |

Dear Editor in chief,

Please find enclosed our manuscript entitled: “**Hippocampal collagen as a potential target for post-surgical treatment; effects of whole-body vibration and exercise**”, which we would like to be considered for publication in Plos One.

A major complication of surgery may be declined mental health, associated with neuroinflammation. Although a derailed inflammatory process as part of wound healing, is indicated to play a major role, anti-inflammatory therapy seems not sufficiently effective to protect neurons from damage. Therefore preservation of neuron integrity during neuroinflammation, by targeting their protective collagen sheet may provide a promising alternative.

Exercise, and the novel therapeutic alternative, Whole Body Vibration (WBV), share combined anti-inflammatory and collagen preserving effects in the peripheral body, but effects in the brain are less known. Aim of the present study was to explore the therapeutic potential of WBV compared to exercise shortly after surgery in rats, with regard to hippocampal neuroinflammation and collagen expression.

WBV as well as exercise restored the surgery-induced declined collagen expression, while partly reversing microglia activation in the CA1 area. Moreover, effects on collagen appeared to be subtype- and region-specific, with overall similar effects of WBV and exercise. Nevertheless, the neuroprotective potential of postoperatively altered brain collagen needs further investigation. Since patients may not capable or unmotivated to benefit from the advantages of physical exercise, WBV may provide a promising alternative.

None of the material presented in the manuscript have been published, nor is under review elsewhere. None of the authors indicated conflict of interest. All authors have read and approved submission of the manuscript.

Hope to hear from you in due course

Yours sincerely,

Dr. Regien G. Schoemaker

Dept. Neurobiology, University of Groningen/

University Medical Centre Groningen

p.a. Nijenborgh 7

9747 AG Groningen

The Netherlands
